## Supplemental Information for "Cell intrinsic dynamics guide neuroblast ingression independent of tissue fluidity"

### Supplementary Figures

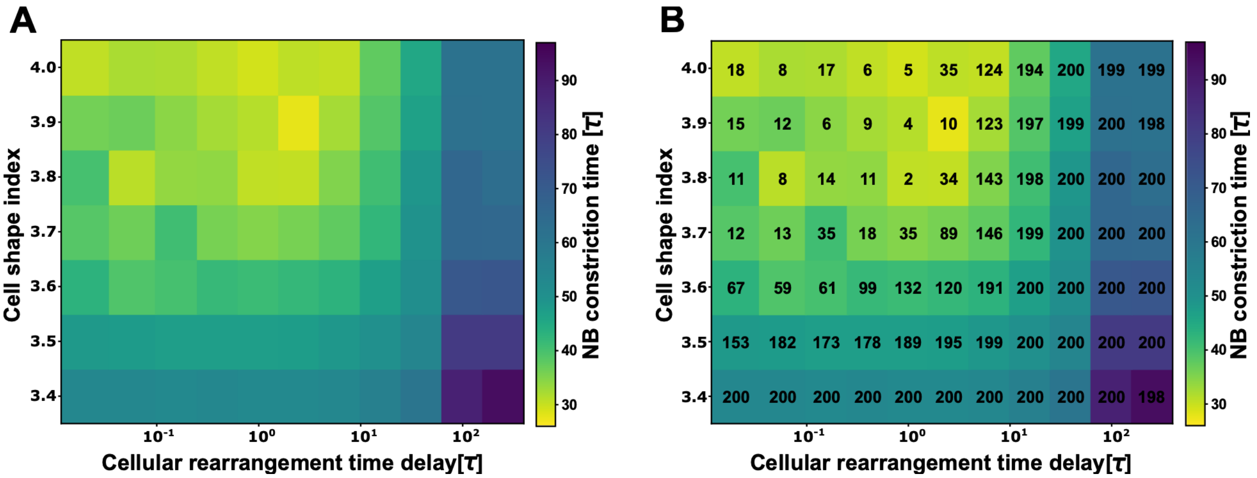


**Figure S1: Combined effects of T1 transition delay and cell shape index on NB ingression timing.** (A) NB constriction time as a function of both T1 cellular rearrangement time (delay) and cell shape index. (B) Same plot as in (A) with the number of successfully ingressed NB cells overlaid in the corresponding parameter grid. Each grid point represents 40 independent simulations with 5 NB cells per simulation (total 200 NB cells per condition). $\gamma_0 = 0.2$, and $\gamma_{ij}^{NB}(t = 0) = 0.1$. Note that missing data occur when either tissue elongation (4-fold) or simulation time limit (200τ) is reached before NB ingression completes, which happens at both extremes of the parameter space: rapid tissue extension (short T1 delay, high shape index) and excessive tissue solidification (long T1 delay, low shape index).

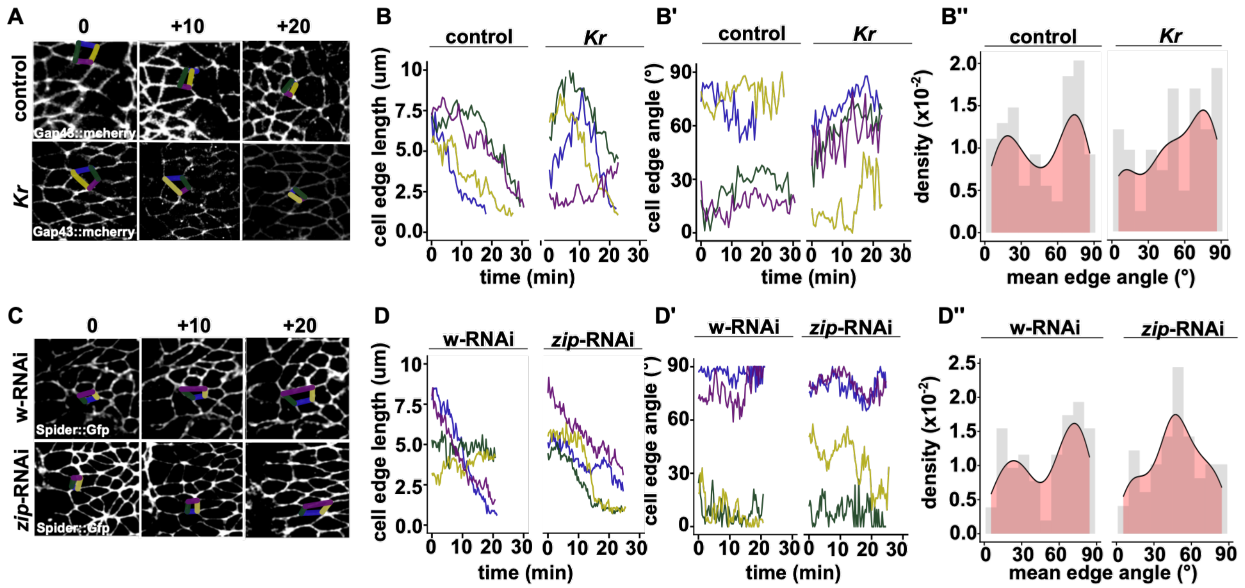


**Figure S2: Loss of anisotropy in *Kr* mutants and *zip*-RNAi.** (A) Representative time-lapse images of ventral epithelium during NB ingression (membrane marker Gap43::mCherry, individual edges color-coded to plots B-B ’). Anterior left, ventral down, t= 0 aligned to start of ingression. (B-B’). Quantification of individual edges from NB tracked in panel A. t= 0 aligned to start of ingression. (B) Edge length, (B’) cell angle. (B”) Probability density distribution of edge angles. Peaks indicate angles with relative higher frequencies (n = 5 embryos, 30 NBs. (C) Representative time-lapse images of ventral epithelium during NB ingression (membrane marker Spider::Gfp, individual edges color-coded to plots D-D ’). Anterior left, ventral down, t= 0 aligned to start of ingression. (D-D’) Quantification of individual edges from NB tracked in panel A. t= 0 aligned to start of ingression. (D) Edge length, (D’) cell angle. (D”) Probability density distribution of edge angles. Peaks indicate angles with relative higher frequencies n = 4 embryos, 20 NBs per genotype.

**Supplementary Movie Captions**

**Movie S1. Simultaneous convergent extension and NB ingression.** Time-lapse visualization of an anisotropic vertex model showing simultaneous tissue convergent extension (cell intercalation driven by anisotropic myosin organization) and NB ingression (apical area constriction). Five ingressing NB cells are embedded within the tissue (highlighted in green). The tissue undergoes planar-polarized cell rearrangements (T1 transitions) characteristic of convergent extension, progressively elongating 2-fold along the anterior-posterior axis. Simulation parameters correspond to control conditions (Methods).

**Movie S2: Neuroblast ingression occurs concurrently with GBE.** Time lapse of PH: mcherry-expressing embryo tracking the apical surface of individual NBs within the ventral column. Video displays 14 frames per second. Time in minutes:seconds.

**Movie S3: Tissue elongation delays in myosin defective embryos.** Time lapse of PH:mcherry embryos in wild-type, *Kr and zip*-RNAi backgrounds. Green line shows tracking of movement of a single cell during GBE. Time is min:sec aligned to the onset of GBE (t=0). Video displays in 7 frames per second.

**Movie S4. Loss of myosin anisotropy delays ingression.** Time-lapse visualization of the model simulating *Kr* mutant conditions with (right) and without (left) loss of NB myosin anisotropy. Tissue undergoes impaired convergent extension with reduced anisotropic line tension (50% of control). Five ingressing NB cells (green) undergo apical constriction with anisotropic tension along AP edges (left) or with isotropic tension applied uniformly to all edges, reflecting the observed loss of myosin polarization in *Kr* mutants (right).

**Movie S5: Loss of anisotropy in *Kr* mutants and *zip*-RNAi.** Time lapse of embryos in wild-type (Gap43::mCherry), *Kr* (Gap43::mCherry) *and* zip-RNAi (Spider::Gfp) backgrounds. Representative NB is segmented in cyan. Time in min:sec. Video displays 7 frames per second.

**Movie S6. *zip*-RNAi simulations: endocytosis-contractility coupling rescues NB ingression in the model despite tissue solidification.** Time-lapse visualization of the model simulating *zip*-RNAi conditions with near-complete tissue solidification. In all panels, tissue undergoes severe reduction in cell rearrangements (10-fold increase in T1 delay) and dramatically reduced anisotropic line tension (25% of control). Five ingressing NB cells (green) undergo apical constriction under three conditions: anisotropic tension along AP edges (left), isotropic tension applied uniformly to all edges (middle), or isotropic tension modulated by endocytic activity via endocytosis-contractility coupling (right). While anisotropic (left) and isotropic (middle) NB constriction both fail to recover normal ingression timing, the coupled endocytosis-contractility model (right) rescues normal ingression despite severe tissue solidification and myosin depletion.
